# MMH: A multimodal dataset of whole-body kinematics, bilateral ground reaction forces, and lower-limb surface electromyography signals during load lifting and lowering

**DOI:** 10.64898/2026.04.22.718747

**Authors:** Mahdi Mohseni, Abdul Aziz Hulleck, Marwan El Rich, Navid Arjmand

## Abstract

This study presents the MMH dataset, a laboratory-collected *in vivo* dataset comprising whole-body kinematics, three-dimensional ground reaction forces and two-dimensional centres of pressure under both feet, as well as surface electromyography (sEMG) signals of twelve lower-limb muscles (six muscles per leg) during load lifting and lowering tasks. Ten healthy, normal-weight, young male adults each performed 72 trials combining one- and two-handed load (2 kg) lifting and lowering. These trials include multiple initial and final load locations while using three different lifting techniques (stoop, semi-squat, and full-squat). The kinematic and force-plate measurements provide rich input for ergonomic risk assessment tools and optimisation-based musculoskeletal models aimed at quantifying and managing musculoskeletal risk of injury. Also, the sEMG recordings enable the development of EMG-assisted musculoskeletal models and support validation of predictions from optimisation-based models. These makes the multimodal MMH dataset a valuable resource for biomechanics, ergonomics, and human movement research.

## 1. Background

Manual material handling (MMH) activities such as load lifting, lowering, carrying, pushing, and pulling are among the most prevalent occupational causes of musculoskeletal disorders (Van Nieuwenhuyse, 2004). Direct *in vivo* measurement of joint loads is invasive, technically demanding and costly (Bender et al., 2024; Dreischarf, Rohlmann, et al., 2016; Wilke et al., 2001). As an alternative, ergonomic risk assessment tools and biomechanical modelling approaches, particularly musculoskeletal models, are widely used to estimate joint loads and muscle forces and to manage injury risk during MMH tasks (Bahramian et al., 2023; Daroudi et al., 2024; Dreischarf, Shirazi-Adl, et al., 2016; Jamshidian et al., 2025; Zargarzadeh et al., 2024).

These tools and modelling approaches typically require as input either whole-body kinematics, ground reaction forces (GRFs), and centres of pressure (CoPs) (for ergonomic risk assessment tools and optimisation-based musculoskeletal models) (Daroudi et al., 2024; Hosseini et al., 2026; Zargarzadeh et al., 2024), or muscle surface electromyography (sEMG) signals (for EMG-assisted models) (Mohammadi et al., 2015; Samadi & Arjmand, 2018). Acquiring such data necessitates specialised and often expensive equipment, including motion capture cameras (Asadi & Arjmand, 2020; Hulleck et al., 2023, 2024), inertial measurement units (Nasrabadi et al., 2022), force plates (Ghasemi & Arjmand, 2021), and sEMG sensors (Gracia-Ibáñez et al., 2024). On the other hand, recent studies have proposed kinematics- and machine learning-based methods to predict body posture/movement (Ahmadi et al., 2025; Hosseini et al., 2025; Hosseini & Arjmand, 2024; Mohseni et al., 2022, 2024), force-plate data (Di Pietro et al., 2025; Fluit et al., 2014; Hosseini et al., 2026), and spinal loads (Arjmand et al., 2013; Hosseini et al., 2026; Mohammadi & Arjmand, 2026).

The present study introduces a new comprehensive public dataset (MMH dataset) comprising whole-body three-dimensional movement data, three-dimensional GRFs and two-dimensional CoP data, and surface EMG signals from 12 lower limbs muscles. These data were collected during a range of load (2 kg) lifting and lowering activities. MMH dataset is intended to support the development of subject specific musculoskeletal models for different body segments, the validation of existing optimisation- and EMG-based models, and future advances in ergonomics risk assessment and MMH biomechanics.

## 2. Methods

### 2.1 Participants

Ten normal-weight young male participants (in mean ± standard deviation format - body mass: 69.5 ± 7.4 kg, height: 179.7 ± 6.7 cm, body mass index (BMI): 21.5 ± 2.3 kg/m^2^, age: 24 ± 2 years) with no history of musculoskeletal disorders in the preceding six months took part in this study. Their age and the following anthropometric data were collected from each participant: 1-body mass and height, 2-segment lengths of the right foot, upper arm, lower arm (forearm), whole leg, upper leg (thigh), lower leg (shank), neck, and shoulder, as well as circumferences of the chest, waist, hip, thigh, ankle, upper arm, and wrist following (Liu et al., 2021), and 3-distances between the right and left anterior superior iliac spine (ASIS) landmarks; distances between ASIS and foot medial malleolus of the foot for both legs; and widths of the right and left knees, ankles, elbows, wrists, hands, and shoulders, according to the Vicon Nexus Plug-in Gait Reference Guide (2023). These anthropometric measurements can be used to post-process the raw kinematic data in various kinematics-based approaches and software environments (Hulleck et al., 2024; Mohseni et al., 2024). The study protocol was approved by the institutional ethics committee (approval ID: IR.IUMS.REC.1401.740), and all participants provided informed consent prior to data collection.

### 2.2 MMH activities

Participants moved a 2 kg load from three ground-level start positions to three chest-level target positions (lifting) and then returned the load to its original position (lowering). The combination of three start and three target locations yielded nine distinct lifting-lowering conditions. At each height (ground- and chest-level), the three positions consisted of one symmetry location in sagittal plane (anterior to the participant) and two asymmetry locations on the right side at 45° and 90° relative to the sagittal plane (Figure 1). For the ground-level positions, the horizontal distance from the load to the participant’s heels were approximately 40 cm. At the chest-level, participants were instructed to hold the load with the arms fully extended, without elbow flexion, at the corresponding symmetric or asymmetric position around their self-selected chest-level vertical position.

**Figure 1:**
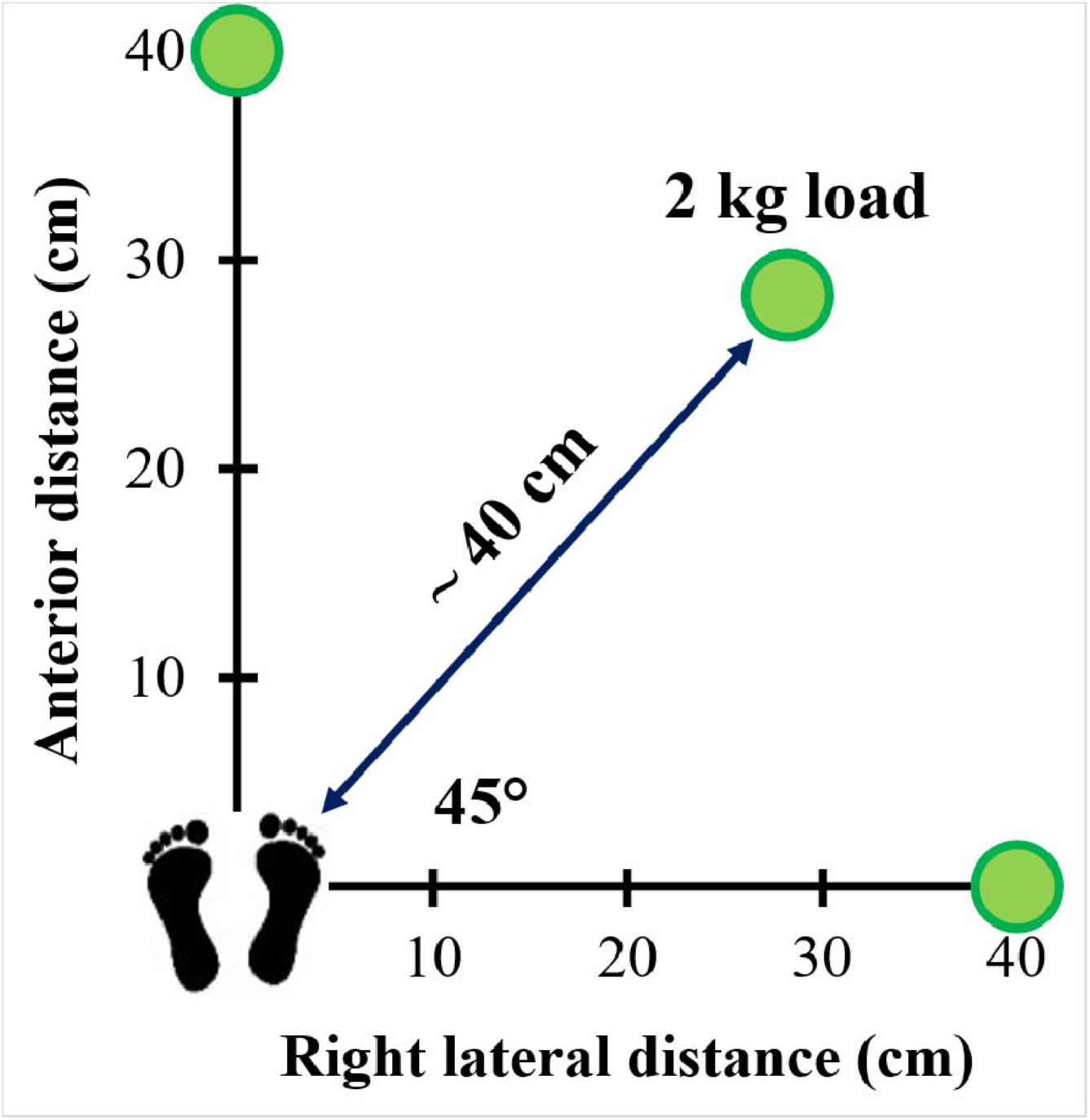
A schematic of ground-level horizontal locations of the load during lifting and lowering tasks.

All tasks were performed using three common lifting techniques: stoop (knees extended, trunk flexed), full-squat (knees flexed, trunk relatively upright), and semi-squat (both knees and trunk flexed) (Mohseni et al., 2024). Stoop tasks were executed with both one-handed and two-handed grips, whereas full-squat and semi-squat tasks were performed only with both hands. All one-handed activities were executed with the right hand. Under this design, each participant completed 36 lifting and 36 lowering trials, for a total of 72 MMH tasks, performed at self-selected load moving speed.

### 2.3 Data collection

During each MMH task, whole-body kinematics were recorded using a ten-cameras Vicon motion capture system (Vicon Motion Systems Inc., Oxford, UK) at 120 Hz. Forty-one passive reflective markers were placed on anatomical landmarks according to an extended Plug-in Gait marker set (Mohseni et al., 2024). The 3D position of the load at its initial and final locations could be approximated from the kinematic data: for one-handed tasks, the load position could be estimated using the 3D coordinates of the right finger marker, whereas for two-handed tasks it could be estimated from the midpoint between the right and left finger markers.

Three-dimensional GRF and two-dimensional CoP data under both feet were measured using two force plates (Kistler Instrument AG, Switzerland) at 1200 Hz. Simultaneously, sEMG signals were acquired bilaterally from tibialis anterior, medial gastrocnemius, lateral gastrocnemius, rectus femoris, semitendinosus, and vastus lateralis (12 muscles in total) using twelve wireless surface EMG sensors (Myon Ltd, Switzerland). sEMG electrode placement followed the recommendations of the sEMG for Non-Invasive Assessment of Muscles (SENIAM) project (Stegeman & Hermens, 2007).

## 3. Data accessibility and potential applications

Following data collection, all motion capture, force-plate, and sEMG data were processed via Vicon Nexus software (v.2.12.1., Vicon Motion Systems Inc., Oxford, UK). Marker trajectories were labelled using the extended Plug-in Gait convention (Mohseni et al., 2024). For each participant, the fully processed datasets from all systems were exported as nine C3D files (L1.c3d to L9.c3d), corresponding to the nine lifting-lowering conditions (Table 1). Each C3D file includes marker trajectories, GRFs, CoPs, and sEMG signals of four lifting and lowering tasks performed while adopting one-handed stoop, two handed stoop, two handed semi-squat, and two handed full-squat lifting techniques, respectively. Also, the corresponding GRF and CoP data are presented in nine MOT files (L1.mot to L9.mot) separately as well. All C3D and MOT files for all the 10 participants of the MMH dataset, together with the associated anthropometric measurements, are publicly available at the following link: https://drive.google.com/file/d/1XfrAZPMvBjeB7FI-ciaypdKjU5urvlHk/view?usp=sharing

**Table 1:** The load location angles related to the sagittal plane in nine output C3D and MOT files.

| C3D-MOT file name | Start load location related to sagittal plane (ground-level) | Target load location related to sagittal plane (chest-level) |
| --- | --- | --- |
| L1 | 0° symmetry | 0° symmetry |
| L2 | 0° symmetry | 45° asymmetry |
| L3 | 0° symmetry | 90° asymmetry |
| L4 | 45° asymmetry | 0° symmetry |
| L5 | 45° asymmetry | 45° asymmetry |
| L6 | 45° asymmetry | 90° asymmetry |
| L7 | 90° asymmetry | 0° symmetry |
| L8 | 90° asymmetry | 45° asymmetry |
| L9 | 90° asymmetry | 90° asymmetry |

The C3D files can be used directly as kinematic and external load (e.g., GRFs and CoPs) inputs in AnyBody Modelling System (AnyBody Technology A/S, Aalborg, Denmark) for the development and evaluation of musculoskeletal models (Bahramian et al., 2023; Daroudi et al., 2024; Dehghan & Arjmand, 2024; Hosseini et al., 2026). An AnyBody full-body musculoskeletal model compatible with the marker configuration used in this dataset is available in can be found in (Andersen et al., 2009, 2010). The same C3D files can also be imported into MATLAB via the BTK Toolkit (Barre & Armand, 2014), which allows conversion of marker and force-plate data to .trc format for use as kinematic file in OpenSim (Daroudi et al., 2024; Dehaghani et al., 2022; Delp et al., 2007; Jamshidian et al., 2025). An OpenSim lumbar spine musculoskeletal model compatible with the present marker set is provided in (Favier et al., 2021). Additionally, the sEMG channels stored in the C3D files can be used for EMG-assisted musculoskeletal modelling, muscle activation pattern and synergy analysis, and for validating muscle force predictions from optimisation-based models, extending the applicability of MMH dataset to a wide range of ergonomic and biomechanical investigations.

## Acknowledgment

We extend our heartfelt gratitude to all the participants for generously volunteering their time to this study. We are also grateful to Mr. Mohammad Madadi and the members of the Djawad Mowafaghian Research Centre for Intelligent Neurorehabilitation Technologies for their invaluable assistance with data collection.

